# Altered Task Demands Lead to a Division of Labor for Sensory and Cognitive Processing in the Middle Temporal Area

**DOI:** 10.1101/2021.12.08.471611

**Authors:** Hayden Scott, Klaus Wimmer, Tatiana Pasternak, Adam C. Snyder

**Affiliations:** Brain and Cognitive Sciences, University of Rochester, Rochester, 14627; USA; Center for Visual Sciences, University of Rochester, Rochester, 14620; USA; Neuroscience, University of Rochester, Rochester, 14620; USA; Centre de Recerca Matem`atica (CRM), Campus de Bellaterra, Edifici C, 08193 Bellaterra, Barcelona, Spain

**Keywords:** Neural Variability, Information, Working memory, MT, primate, vision

## Abstract

Neurons in the primate Middle Temporal (MT) area signal information about visual motion and work together with the lateral prefrontal cortex (LPFC) to support memory-guided comparisons of visual motion direction. These areas are reciprocally connected, and both contain neurons that signal visual motion direction in the strength of their responses. Previously, LPFC was shown to display marked changes in stimulus coding with altered task demands, including changes in selectivity for motion direction, trial-to-trial variability in responses, and comparison effects. Since MT and LPFC are directly interconnected, we sought to determine if MT neurons display similar dependence on task demands. We found that active participation in a motion direction comparison task affected both sensory and non-sensory activity in MT neurons. In fact, neurons that became less-selective for motion direction during the active task showed increased signalling for cognitive aspects of the task. This heterogeneity in neural modification with heightened task demands suggests a division of labor in MT, whereby sensory and cognitive signals are both heightened in different subpopulations of neurons.

## Introduction

The Middle Temporal (MT) area of monkey cerebral cortex is a well-studied extrastriate visual area known to have robust selectivity for visual motion features (Britten et al. 1992; Born and Bradley 2005; Pasternak and Tadin 2020). Motion signals represented by the activity of neurons in this area are likely to contribute to memory-guided comparisons of visual motion direction (Lui and Pasternak 2011; Bisley, Zaksas, and Pasternak 2001; Katz et al. 2016). Lesions of this area impair the ability to perform direction discrimination during motion comparison tasks (Rudolph and Pasternak 1999; Bisley and Pasternak 2000); whereas electrical stimulation of this area can bias the behavior of an animal as though a particular direction of motion was seen that was not present in the stimulus (Bisley, Zaksas, and Pasternak 2001; Salzman et al. 1992; Salzman, Britten, and Newsome 1990). These findings indicate a key role for MT in representing behaviorally relevant visual motion information and demonstrate that MT has an important role in task-related perceptual decisions. One open question, however, is how behavioral goals impact the neural activity in area MT during sensory (e.g., encoding/decoding of stimulus) and non-sensory (e.g., reward contingencies, decision making, etc.) periods of the task.

It is likely that during memory-guided comparisons of motion direction, area MT acts in concert with another brain area, the lateral prefrontal cortex (LPFC). LPFC and MT are reciprocally connected (Ungerleider and Desimone 1986; Barbas 1988; Petrides and Pandya 2006), and both are involved in memory-guided motion comparison tasks (Zaksas and Pasternak 2006). Neurons in LPFC contain direction information during motion discrimination tasks, but this selectivity is largely attenuated when animals do not have to make perceptual decisions, or when that information is not relevant to the task (Hussar and Pasternak 2009; Hussar and Pasternak 2012; Hussar and Pasternak 2013). Previous research has also found reduced trial-to-trial variability in LPFC with increased task demands (Hussar and Pasternak 2010), suggesting LPFC neurons may be recruited to support task performance when needed. Because of the interconnected structural and functional relationship between LPFC and MT, such changes in LPFC activity with task demands may be reflected in MT dynamics and altered processing of stimulus information, but this hypothesis remains untested.

To investigate this question, we measured how activity in MT was affected by different demands encountered during an active task that required monkeys to report memory-guided comparisons of motion direction, and a passive, identically structured task involving the same stimuli that did not require the animals to report their decisions. The active task consisted of two random dot motion stimuli separated by a delay, where monkeys were rewarded for correctly reporting whether or not the direction of motion in the second stimulus (“S2”) matched the first (“S1”). During the passive task, signaled by a distinct fixation target, the trial structure and sensory conditions were the same, but required no perceptual decision, and the reward was delivered on each trial. Thus, despite identical sensory conditions, the task demands during active and passive tasks were different; the active task can be parsed as a sequence of: stimulus encoding (S1), memory maintenance (delay period), recall and comparison (S2), and finally perceptual report (S2/post-S2). We hypothesized that the sensory responses to motion in S1 and S2 during the active task would be reflected in sharper tuning, reduced trial-to-trial variability, and/or modulation gain, in line with prior findings about selective attention to motion direction (Ponce-Alvarez et al. 2013; Cohen and Newsome 2008; Arandia-Romero et al. 2016).

We found that the effects of task demands depended on the amount of task-relevant stimulus information (e.g. direction selectivity) neurons conveyed. Specifically, we found three main modulation profiles for stimulus processing in the active task compared to the passive task: neurons that showed increased direction information (“IDI” neurons), neurons that showed decreased direction information (“DDI” neurons), and neurons with with largely the same amount of direction information during the two tasks (“SDI” neurons). These profiles likely reflect neural heterogeneity and should therefore be taken as soft boundaries more-so than distinct groupings. We found that during the active task, IDI neurons displayed tuning gain and a reduction in trial-to-trial variability, but showed similar comparison signals during the two tasks. In contrast, the task effects for DDI neurons were reflected in weaker tuning for direction, but not in trial-to-trial variability. The task effects on direction tuning and response variability of the SDI neurons were weak. Instead, these neurons, similarly to the DDI neurons, showed task dependent comparison signals. These results suggest a division of labor in area MT, grading sensory processing and cognitive effects of the task across subpopulations of neurons.

## Methods

### Resource Availability

#### Lead Contact

Requests for further information should be directed to the lead contact Adam Snyder.

#### Materials Availability

This study did not generate any new reagents.

#### Data and Code Availability

Data and code used for this paper are made available by request through the lead contact.

### Subjects

Subjects used in this study were three adult male rhesus macaques m201, m202, and m317. All training, surgery, and experimental procedures were performed in accordance with the National Institutes of Health *Guide for the Care and Use of Laboratory Animals* and were approved by the University of Rochester Committee for Animal Research. Surgery was performed using aseptic technique with isofluorane general anesthesia and perioperative opiate analgesics. An initial surgery was performed to implant a surgical steel head holder restraint embedded in bone cement secured to the cranium with ceramic bone screws. After behavioral training, a subsequent surgery was performed to implant a PEAK canula (19 mm diameter; Crist Instruments, Hagerstown, MD) over MT.

### Data Acquisition

Recording chambers enclosed the craniotomies and contained 1mm-spaced CILUX grids (Crist Instruments). Thirty-two-channel S-probes (Plexon, Dallas Texas) were lowered through custom-made steel guide tubes, which were themselves inserted just low enough to penetrate the dura. We then drove the electrodes using a NAN electrode drive (NAN Instruments, Nof Hagalil, Israel). Recordings were done by coordinating a Plexon (Dallas, Texas) Multichannel Acquisition Processor and the data acquisition system TEMPO (Reflective Computing).

### Receptive Field Mapping

Prior to each recording session, receptive fields were mapped in order to identify where the stimuli should be displayed. The initial manual phase consisted of moving a patch of dot motion around on the stimulus display with a computer mouse while listening to spiking activity via an audio monitor. Afterwards, we presented small dot motion patches (1°) in different directions on the vertices of a grid spanning the likely RF area identified in the manual step. Stimuli for each session were placed such that they covered the aggregate receptive field area for the recording (Figure 1C).

**Figure 1.**
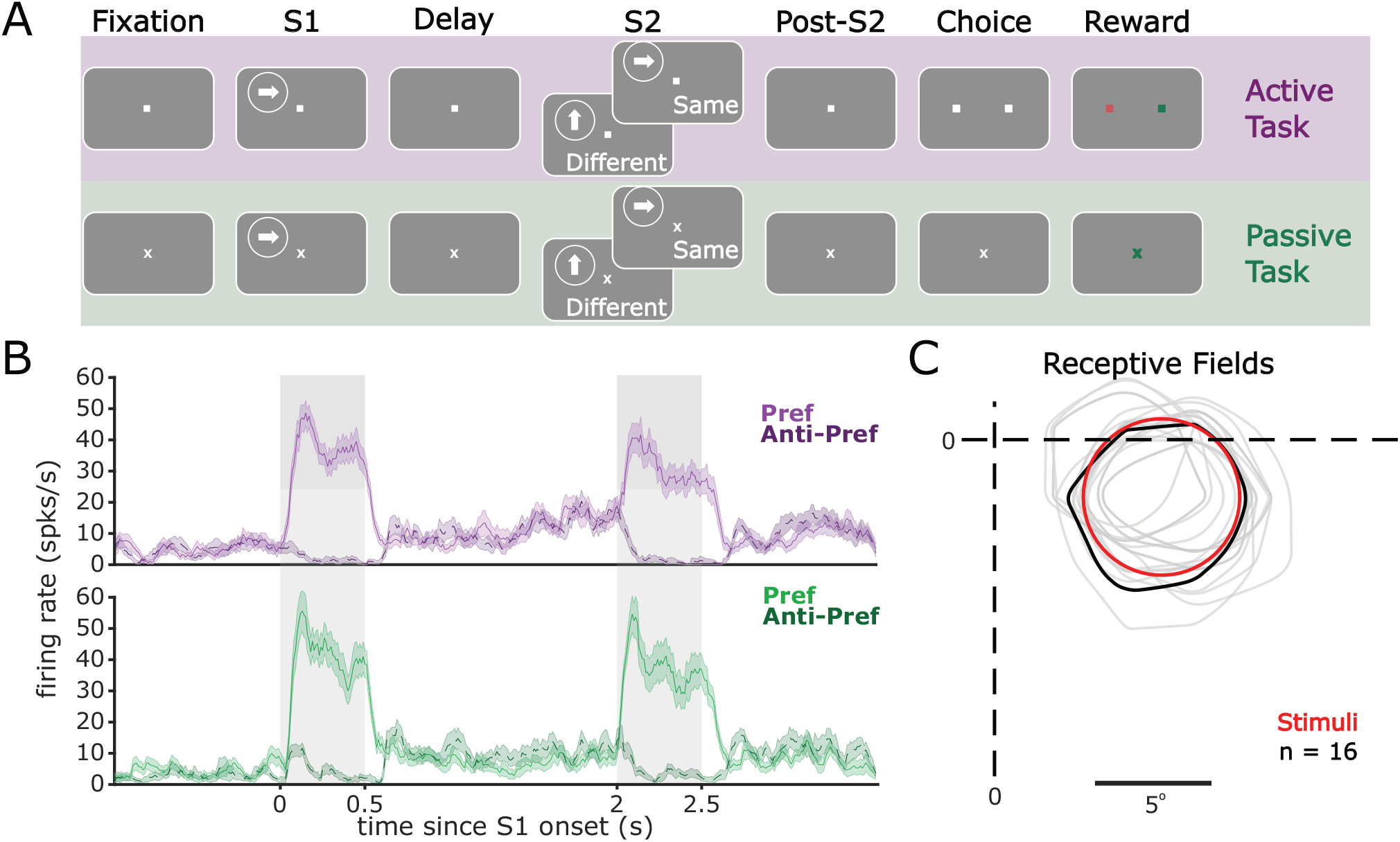
Task design and task modulation of MT firing rates. A.) Trials consisted of a pre-stimulus 1 second fixation period, a first stimulus (S1), followed by a 1.5 second delay, then a second stimulus (S2) and a post-S2 fixation period. After the 1 second post-S2 fixation period subjects were either rewarded (passive task) or had to report a decision with a saccade (active task). During the active task, correct choices were rewarded with juice while incorrect choices resulted in a 3 s time-out signaled by a tone and no reward. During S1, stimuli moved in one of 8 directions (0°, 45°, 90°, 135°, 180°, 225°, 270°, 315°), followed by the S2 that moved either in the same direction, or 90°off of S1 (rotated left or right). B.) PSTHs of an example neuron recorded in both tasks. Solid line represents the neuron’s response to its preferred direction while dashed lines indicate 180∘ away from preferred (“anti-preferred”). C.) Receptive fields (grey) of each simultaneously recorded neuron from one example session, along with the location and size of the stimuli (red). Black curve corresponds to example neuron in B. Contours represent isointensity contours at 50% of the peak response.

### Stimulus Presentation

The stimuli and the behavioral tasks were similar to those used in previous studies (Hussar and Pasternak 2012; Wimmer et al. 2016). Stimuli were presented at 75 Hz monitor refresh rate on a 19-inch IIyama Vision Master Pro-513 monitor with 1,152 by 870 pixel resolution. The stimuli consisted of random dots placed in a circular aperture fit to maximize the overlap of the recorded receptive fields. The dot density was 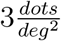 and each dot was 0.03 degrees of visual angle (DVA) in diameter with a luminance of 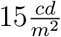 shown on a dark background of 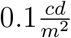. The stimuli moved in one of 8 directions (0°, 45°, 90°, 135°, 180°, 225°, 270°, 315°), and their sizes ranged from 2 DVA to 15 DVA in diameter depending on eccentricity. Monkeys were required to maintain fixation within 1.5 of a central dot throughout the trial until the response interval, and eye position was monitored with an ISCAN infrared eye-tracking device.

### Task Design

Trials started once subjects fixated on a central dot. They then had to maintain fixation for 1 second before S1 onset. Both S1 and S2 lasted 500ms. S1 was followed by a 1.5 second delay period, and then the second stimulus S2 was presented. Subjects then had to wait an additional 1 second after S2 offset before either a choice was required (active task), or they were simply rewarded (passive task). For the active task, the central fixation point was extinguished at the beginning of the response interval, and two identical choice targets were presented 5 DVA to the right and left of fixation, and the animal indicated its judgement as to whether S1 and S2 were the same direction or not by making a saccade to the right or left choice target, respectively. In most recording sessions, we collected data for both the active and the passive tasks. We always collected data for the active task before the passive task, because animals would have been unmotivated to perform the more difficult active task if they had performed the easier passive task first.

### Data Analysis

Spike sorting was done offline with Plexon sorter (Version 3.3.2) by using principal component analysis (PCA) on the spike waveforms and manually clustering. A neuron was included for analyses if there were at least ten trials in each direction condition, and if the stimuli covered at least 30% of its 50% isointensity response field. A total of 1264 neurons across both tasks passed these criteria (m201: n = 263, m202: n = 454, m317: n = 547), with 928 neurons recorded in the active task and 336 neurons recorded in the passive task. Of those neurons, 254 were confidently matched across task and included for further analysis. Neurons were considered matched across task if they were recorded on the same channel and their waveforms were highly correlated (*r*^2^ > 0.99). All subsequent analyses were done using custom routines for Matlab 2020a (The MathWorks, Inc, Natick, MA).

### Mutual Information

Mutual information (MI) quantifies information (in bits) shared between two variables, i.e.: how much the uncertainty about one variable (motion direction condition, in our case) may be reduced with knowledge about the second variable (spike counts). MI provides a robust metric of direction selectivity that accounts for changes in tuning curves, variability, and absolute firing rates. We calculated MI using:

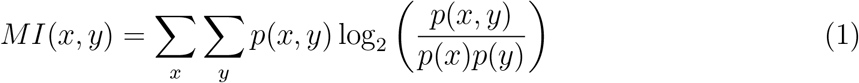

Where *p*(*x, y*) is the joint probability distribution between number of spikes and direction condition, and *p*(*x*)*p*(*y*) is the product of marginal probability distributions. This provides how many bits of information a neuron’s firing rate contains about direction and vice versa. For *p*(*x*), we used the probability of observing a spike count in a bin of width Δ*x* sp/s, where the bin width was determined by “Scott’s rule” (Scott 1979):

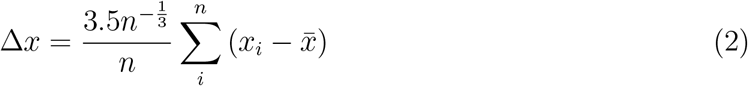

We calculated the mutual information between spike count and S1 direction for each non-overlapping 50ms bin from the start of each trial up to the end of the delay, and then the MI between spike count and S2 direction through the end of the trial. In order to correct for spurious effects that come from differences in trial-counts, we baseline-corrected MI by calculating it many times with shuffled conditions, and subtracted this out (Hatsopoulos et al. 1998).

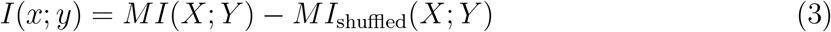

### Classification of neurons into information Enhanced, Suppressed, and Consistent subgroups

We calculated baseline-corrected mutual information (MI) between the firing rate of a neuron and motion direction shown for each time bin. From trial start up to S2 onset MI was calculated according to S1 direction. From S2 onset through the end of the trial, it was calculated between firing rate and S2 direction.

We then took the MI time series for a neuron in active, minus passive, in order to get a task-effect of motion information over time. Note that this was only possible for neurons that were successfully matched across tasks (n=254). The clustering algorithm contains 4 distinct steps.

1. Calculate the Pearson distance (i.e., one minus the Pearson correlation divided by 2) between each pair of neurons’ task-effect time series. This was done on a window during S1. We did this in order to disentangle the sensory and cognitive components, since clustering on S2 would create a circularity for later analyses.
2. Reduce the dimensionality with multidimensional scaling to five dimensions (to mitigate the “curse of dimensionality” for clustering with high-dimensional data).
3. Model selection to pick the appropriate number of clusters. This consisted of fitting multiple Gaussian mixture models to the data, each with different numbers of components (using the MATLAB ‘gmdistribution’ class). We then selected the appropriate number of clusters by selecting the model with the lowest Akaike Information Criterion (AIC), which was three clusters.
4. K-means clustering using the appropriate number of components (3) determined in step 3.

In this way, neurons were partitioned into groups based on how motion information signaling changed across the active and passive tasks. This allows us to then see how other task-related effects may depend on the type of information modulation a neuron receives.

### Resampling Procedure for Clustering Robustness

In order to confirm the validity of our clustering approach, we used a resampling procedure with 100 iterations. We calculated mutual information for each neuron, using random samples of 50% of trials (on each iteration, we used the remaining half of trials to analyze changes in tuning curves; see section *Tuning Curves*). We then ran the clustering algorithm on each iteration, and calculated the most parsimonious model each time using AIC. The majority of iterations resulted in 3 clusters of enhanced, suppressed, and consistent direction information, while no iterations resulted in a single cluster being the best model (Supplemental Figure 1).

### Entropy Factor

Entropy Factor (HF) quantifies the entropy of a neuron’s spike count (across trials) relative to a Poisson process with the same rate parameter,,, *\* (Rajdl, Lansky, and Kostal 2017). It is similar to Fano factor, in that a value of 1 indicates equivalence to a Poisson process, while values greater or less than 1 indicate distributions that are supra-poisson or sub-poisson, respectively. However, unlike Fano factor, which compares the variance of a distribution of spike counts to its mean, entropy factor takes into account the entire distribution of spike counts. This provides a more reliable metric than Fano Factor, particularly when there are a low number of observations (Supplemental Figure 2). For each 50ms time bin, *t*, we calculated the entropy 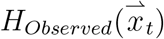 of the distribution of spike counts 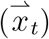 across trials with the same stimulus direction and divided it by the entropy of a simulated Poisson distribution with the same intensity and number of trials.

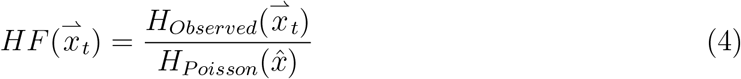

Here, 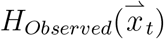 is the entropy of the distribution of spike counts across trials *i* ∈ [1, *n*] for a neuron at time *t*, defined by:

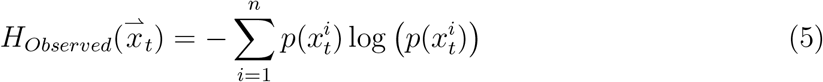

where 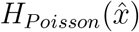 is the entropy of a simulated Poisson process with rate parameter

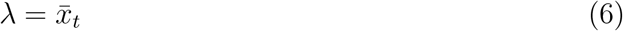

The entropy of a Poisson process was calculated by doing 200 simulations of spike counts for *n* trials pulled from a Poisson distribution with rate parameter,, *\*_*t*_, and the same unbiased bins used for the data (eq. 2). By then averaging the observed entropies of the simulations, we get an estimate of how variable, on average, we would expect the matched Poisson process to be. This normalization accounts for bias in sample entropy due to differences in firing rate or trial count.

### Tuning Curves

Tuning curves were calculated as the average firing rate during the stimulus window for each of the 8 directions. For analyzing tuning curves, we used the half of trials that was not used for clustering based on mutual information from each resampling iteration (see section *Resampling Procedure for Clustering Robustness*), and averaged results across iterations. For testing the effect of stimulus (S1 vs. S2) on tuning and HF, we used a three-way ANOVA with factors of ‘direction’ (eight levels corresponding to each motion direction), ‘stimulus’ (two levels: S1 vs. S2) and ‘task’ (two levels: active vs. passive).

To test whether changes in tuning functions could be explained through changes in amplitude, width, preferred direction, or baseline shifts, we fit Gaussian functions to the responses of individual neurons using the trust-regions algorithm implemented by the Matlab ‘cfit’ object class. The fitted function was

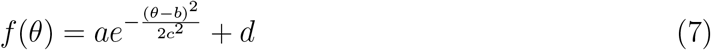

where *a* is the amplitude (in spikes/s), *b* is preferred direction (in degrees), *c* is the width parameter in degrees —related to the full width at half height (FWHH) by *FWHH* = 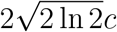, *d* is the baseline response (in spikes/s), and *θ* is the motion direction of a stimulus. Because motion direction has a periodic domain ([0, 360) degrees) whereas the Gaussian function does not, we “centered” the responses of each neuron so its preferred direction of motion was around zero prior to fitting, then we applied the inverse shift to the estimated *b* parameter. We imposed the following constraints: 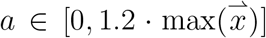, *b* ∈ [−45, 45], *c* ∈ [1, 90], 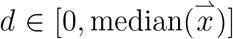, where 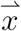 is the set of observed responses (average firing rate during stimulus window) to be fit for a given neuron. The starting point for fitting was 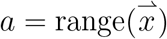, *b* = 0, *c* = 40, 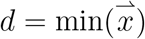.

To test for differences in fitted parameters between tasks, we performed paired t-tests with *alpha* = 0.05, Bonferroni-corrected for the three groups of neurons tested (i.e., DDI, IDI, and SDI). We report the results of this analysis using all neurons, but we also tested subsamples of neurons restricted to have *R*^2^ values for their fits greater than 0.2 (68% of neurons) or 0.5 (34% of neurons) and found that the pattern of results was similar to the results when testing the full sample, and the conclusions were not affected. Note that we used individual trials rather than within-condition averages for fitting tuning functions, so some of the variance unexplained by the fitted model includes trial-to-trial variability.

For reports directly comparing MI to tuning parameters, we used the above procedure on the half of trials not used to calculate MI across all 100 cross-validation folds. This allowed us to avoid any circularity in our comparison of tuning and mutual information.

### Comparison Effects

Comparison effects (CE) were calculated as the area under the curve (AUC) of the receiver operating characteristic. For each neuron, trials were separated by direction condition (during S2), and then by trial type (same direction or different direction trials depending on whether S1 matched S2). CE was calculated on firing rates in 100 ms bins, sliding by 10 ms steps, as the AUC between ‘same’ trials and ‘different’ trials within direction. AUC values were then averaged across their preferred direction and ±45° off preferred within each time bin, in order to reduce noise in the estimate (see Supplemental Figure 4 for a comparison of CEs to preferred versus anti-preferred directions). This resulted in a time course of AUC values for each neuron where values greater than 0.5 indicates a neuron that responds more robustly to identical visual stimuli on trials where the directions of S1 and S2 matched (*S* > *D* neurons), while values less than 0.5 indicated the opposite (*D* > *S* neurons).

### Statistical Significance of CE

For testing task effects on the population average CE (Figure 5), we first averaged for each neuron within each of two time windows of interest: (1) the period when S2 was presented, and (2) the period from S2 offset to the onset of the choice targets. We then tested for a difference between tasks using a paired t-test, with each neuron as a degree of freedom (*α* = 0.01, two-tailed).

Statistical significance of CE for individual neurons (Figure 6) was determined using a bootstrapping procedure. Trial labels (‘same’ or ‘different’ trial) were randomly permuted, and AUC values were calculated 5000 times per neuron following the procedure described above (section *Comparison Effects*), and then averaged within each time period of interest (S2, and post-S2). A neuron was considered significantly CE signaling (Figure 6) if its AUC was significantly different from the bootstrap distribution (*α* = 0.01, two-tailed). Observed CE values greater than the bootstrap distribution indicate same-preferring (*S* > *D*) and lesser than the bootstrap distribution indicate different-preferring (*D* > *S*).

## Results

Three adult male Rhesus macaques (*Macaca mulatta)* performed delayed visual motion comparison tasks while we recorded neural activity in MT (Figure 1A). In the active task, monkeys indicated whether a second stimulus (“S2”) moved in the same or an orthogonal direction to a first stimulus (“S1”) by making a saccade to one of two choice dots. All three monkeys performed above chance in the active task (m201, 10 sessions: 77.79% ± 1.08% accuracy, m202, 28 sessions: 81.63% ± 0.95%, m317, 16 sessions: 85.77% ± 1.36%; *mean* ± *Std*). The passive task, except for a different fixation point shape, was identical up to the choice window, at which time no choice target stimuli were shown, and no further action was required to receive reward (Figure 1A). Note that during both tasks, the timing of salient events, i.e., the onset time of S1 and S2 during both tasks was highly predictable, since on each trial the periods preceding S1 (1 s fixation period) and S2 (1.5 s delay) were always the same. Similarly, on each trial there was always a 1 s period separating the offset of S2 and the decision/reward (active task) or the reward (passive task). We were interested in how neurons’ signaling for motion direction changed from the passive to the active task, and how that related to changes in neurons’ signaling for cognitive processes. The cognitive signals we considered were anticipatory ramping of firing rates and comparison effects. We were unable to consider another cognitive signal of potential interest, choice probability (Britten et al. 1996), because error trials were relatively infrequent in the active task, and no choices were made in the passive task.

Neurons were recorded between 2 and 25 eccentricity, with receptive field sizes proportional to distance from the fovea. A linear fit to receptive field size vs. eccentricity resulted in an intercept of 2.37 and a slope of 0.44, comparable with previous results from this lab (Bisley et al. 2004) and other studies in MT (Albright and Desimone 1987). Neurons were included for analysis if: 1.) Their 50% isointensity response field (see Methods) was at least 30% covered by the stimuli, 2.) the subject did at least 10 trials in each direction condition, and 3.) they were confidently recorded across both tasks (see Methods). This resulted in 254 well-defined neurons included for analysis.

We were interested in the impact of cognitive demands on MT activity. During the active task, such demands varied over the course of a trial, including periods where stimuli were present (i.e., during S1 and S2) and also periods where stimuli were absent (i.e., pre-S1, the delay, and post-S2). For example, during the 1 second fixation period prior to S1, the animal is likely to prepare cognitive resources in anticipation of the stimulus, the appearance of which was highly predictable. Then, during S1, the animal must process and store task-relevant information (motion direction). During the delay, this information must then be maintained in working memory and the appearance of S2 may be anticipated. During S2, task-relevant information must again be encoded, the stored information about S1 retrieved, and a comparison judgement made. These dynamic task demands may result in varying effects on MT activity across these different stages. We first describe the effect of task demands on signaling motion direction by MT activity during S1 and S2. Second, we will describe the effects of task demands on MT activity when stimuli were absent. Third, we will describe effects of task demands on MT activity during the comparison stage of the task, i.e., during the S2 and the post-S2 period preceding the decision/reward (active task) and the reward (passive task).

### Task effects in MT depend on neural tuning

We hypothesized that the accuracy of encoding of motion direction would improve when that information was task-relevant (i.e., in the active task compared to the passive task). To test this, we measured the mutual information (MI) between neural firing rates and the motion direction of S1 and S2. MI quantifies how much information about one signal is provided by another signal (and vice-versa) and takes into account both the tuning of neural responses with respect to a stimulus as well as trial-to-trial response variability (Hatsopoulos et al. 1998; Quian Quiroga and Panzeri 2009).

Task-dependent selectivity would be consistent with the role of MT as a mid-level sensory area. That is, switching from passive viewing to a task requiring judgment of motion direction might have a particularly strong effect on neurons strongly signaling motion direction compared to neurons that did not signal that information so strongly. To test this hypothesis, we calculated the task effect of motion information for each neuron by subtracting the MI during the passive task from the active task (Figure 2 for examples). This approach provided a metric which quantifies, at each time point, how each neuron signals motion direction information during the two tasks (Figure 2D). We then calculated the Pearson distance between each neuron, which provided a dissimilarity matrix for our population of neurons. Neurons close together in this space have similar task effects, whereas neurons far apart are different (Figure 2E). We then reduced the dimensionality to D=5 with multidimensional scaling to minimize distortion of the relative Pearson distances and clustered neurons using a Gaussian Mixture model. We fit models with 1 to 5 clusters using 50% trial hold-out cross-validation. We then took the median Akaike Information Criterion (AIC) across folds and found that three clusters of task effects represent the most parsimonious model (1 Cluster: *AIC* = 6.655 · 10^3^, 2 Clusters: *AIC* = 6.626 · 10^3^, 3 Clusters: *AIC* = 6.624 · 10^3^, 4 Clusters: *AIC* = 6.630 · 10^3^, 5 Clusters: *AIC* = 6.638 · 10^3^; smaller values indicated better models). Importantly, the model with a single cluster was never the model with the best AIC on any cross-validation fold, suggesting that the presence of cluster structure in the data is robust. Nevertheless, we caution that cluster identities are best conceived as “soft” boundaries, and interpret our results as reflecting heterogenous variation across the population. We label the task effects corresponding to these clusters as: Increased Direction Information (IDI), Decreased Direction Information (DDI), and Similar Direction Information (SDI) groups (Figure 3A&B), referring to the average task effect on direction information for neurons within each cluster.

**Figure 2.**
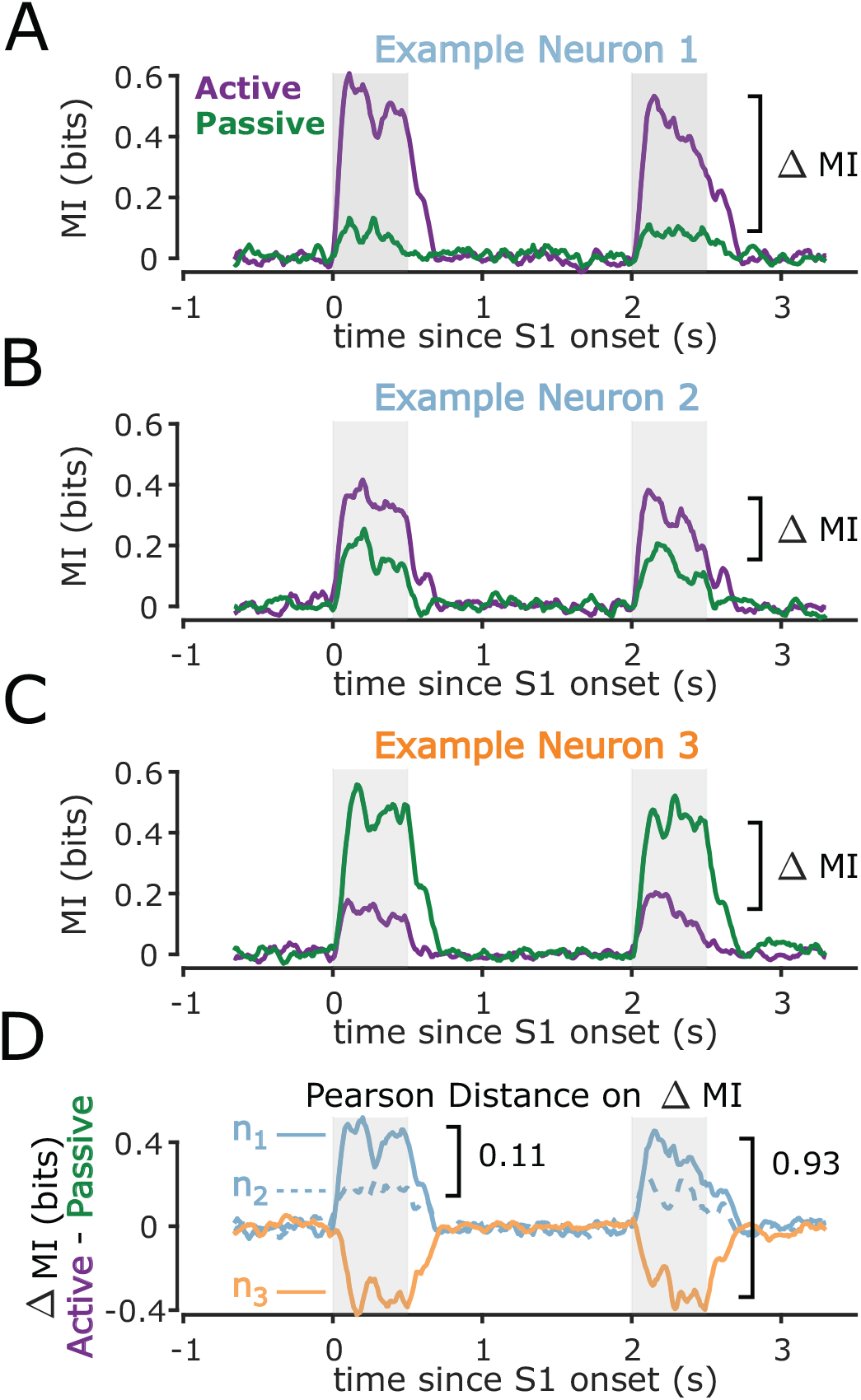
Example Mutual Information Time-Courses. A.) Mutual information between motion direction and firing rate for an example neuron in the active and passive tasks. B-C.) Same as A for two more example neurons. D.) Task effect on motion information (i.e., MI during active minus MI during passive) during the course of the trial for the three example neurons. Pearson Distance was calculated on these time courses for each pair of neurons. Distance values close to 0 mean similar task effects, whereas distance values close to 1 indicate opposite task effects. We then used the set of pair-wise Pearson distances to cluster neurons based on their task effects for MI.

**Figure 3.**
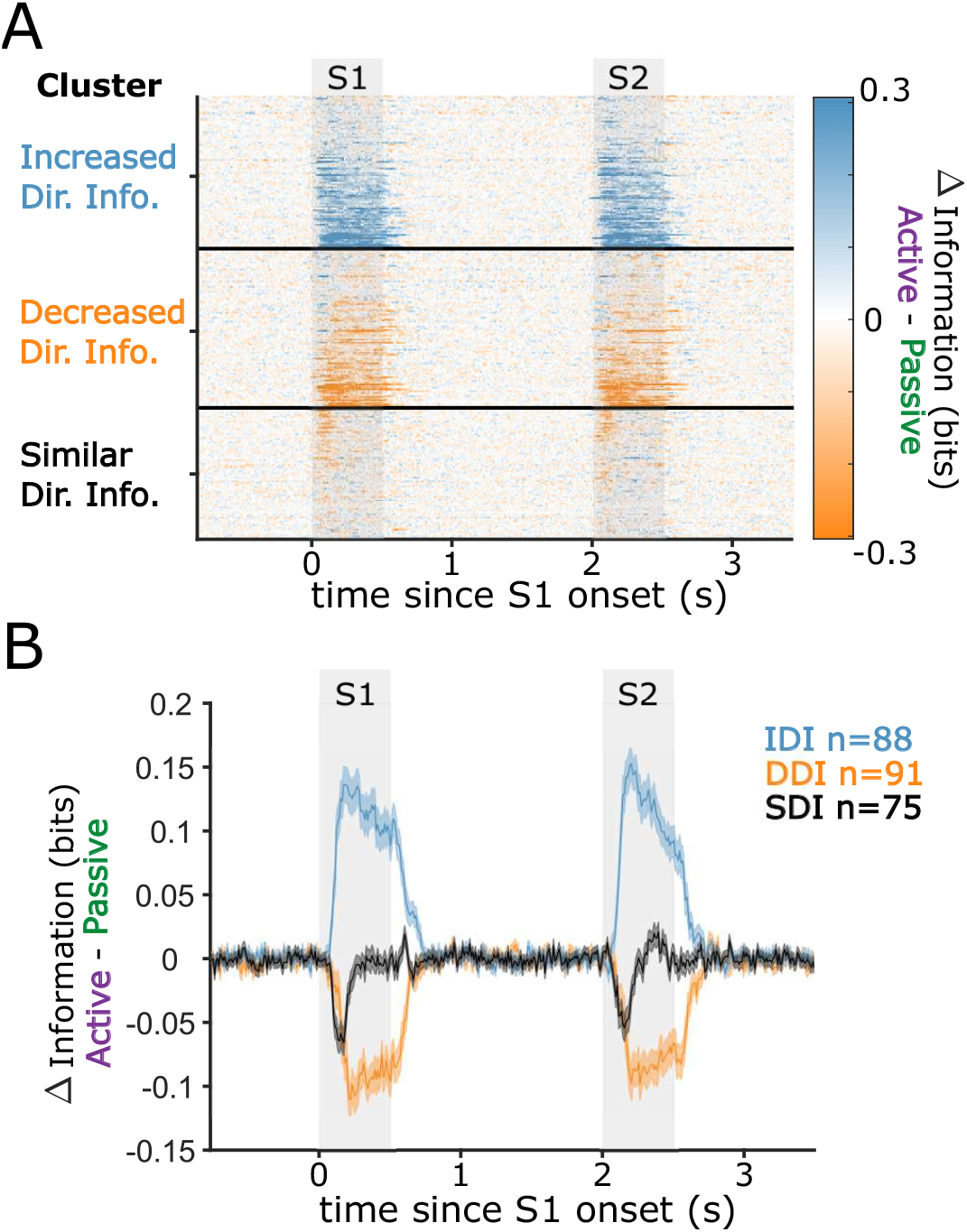
Motion Information Clustering. A.) Time series for active minus passive motion information (MI) for each neuron. Rows correspond to neurons and color represents the difference in information. See Methods for the clustering algorithm. B.) Within-cluster averages from panel A.

Remarkably, during the active task a substantial group of neurons displayed a marked reduction in MI (DDI neurons), indicative of a detriment in direction coding, while the IDI neurons followed the expected pattern of improved direction signaling. These results suggest that heightened task demands result in task-relevant information becoming disproportionately represented in subgroups of neurons.

### Differential Effect of Task Demands on Tuning for Motion Direction

We found that mutual information between neuron firing rates and motion direction changed when motion direction was task-relevant (active task) compared to when it was not (passive task). Since both the tuning for motion direction and trial-to-trial variability contribute to MI, the observed changes in MI could be consistent with a change in tuning, a change in trial-to-trial variability, or a mixture of both. To better characterize the nature of these effects, we separately analyzed tuning and trial-to-trial variability for IDI, DDI, and SDI neuron groups.

We tested for differences in responses to motion for each group by calculating the average firing rate for each neuron, and compared effects of task, direction condition, and stimulus epoch with 3-way ANOVAs. The main effect of ‘stimulus epoch’ (S1 vs S2) was negligible for each group (IDI: *p* = 0.238; DDI: *p* = 0.187; SDI: *p* = 0.531), so for visualization purposes we plotted S1, which unlike S2 does not involve concurrent comparison processes (Figure 4A-C). As expected, all three groups were significantly tuned to motion direction (3-way ANOVA: IDI: *F* = 96.09,*p* = 5.47 · 10^−126^; DDI: *F* = 107.77,*p* = 1.49 · 10^−140^; SDI: *F* = 49.72,*p* = 2.70 · 10^−66^). We found main effects of task in all three groups (IDI: *F* = 15.27,*p* = 9.54 · 10^−5^; DDI: *F* = 18.28,*p* = 1.97 · 10^−5^; SDI: *F* = 7.96,*p* = 0.005), indicating slight changes in overall firing rates with task condition (IDI, active: 27.36 ± 0.52 sp/s mean ± SEM, passive: 24.96 ± 0.44 sp/s; DDI, active: 25.47 ± 0.41 sp/s, passive: 27.82 ± 0.46 sp/s; SDI, active: 23.60 ± 0.43 sp/s, passive: 22.12 ± 0.37 sp/s). We also found interaction effects between task and stimulus direction for IDI and DDI neurons, indicative of task-dependence changes in direction tuning. Firing rates around the preferred direction became enhanced in IDI neurons in the active task (*F* = 5.25,*p* = 5.76 · 10^−6^: Figure 4A,K). In contrast, in the DDI group, firing rates around the preferred direction were reduced in the active task compared to the passive condition (*F* = 4.24,*p* = 1.16 · 10^−4^ Figure 4B,K). The SDI group had no such interaction effect (*F* = 0.24,*p* = 0.974). Taken together, these results imply that direction tuning differs between tasks for IDI and DDI neurons, while task demands do not alter direction tuning of SDI groups.

**Figure 4.**
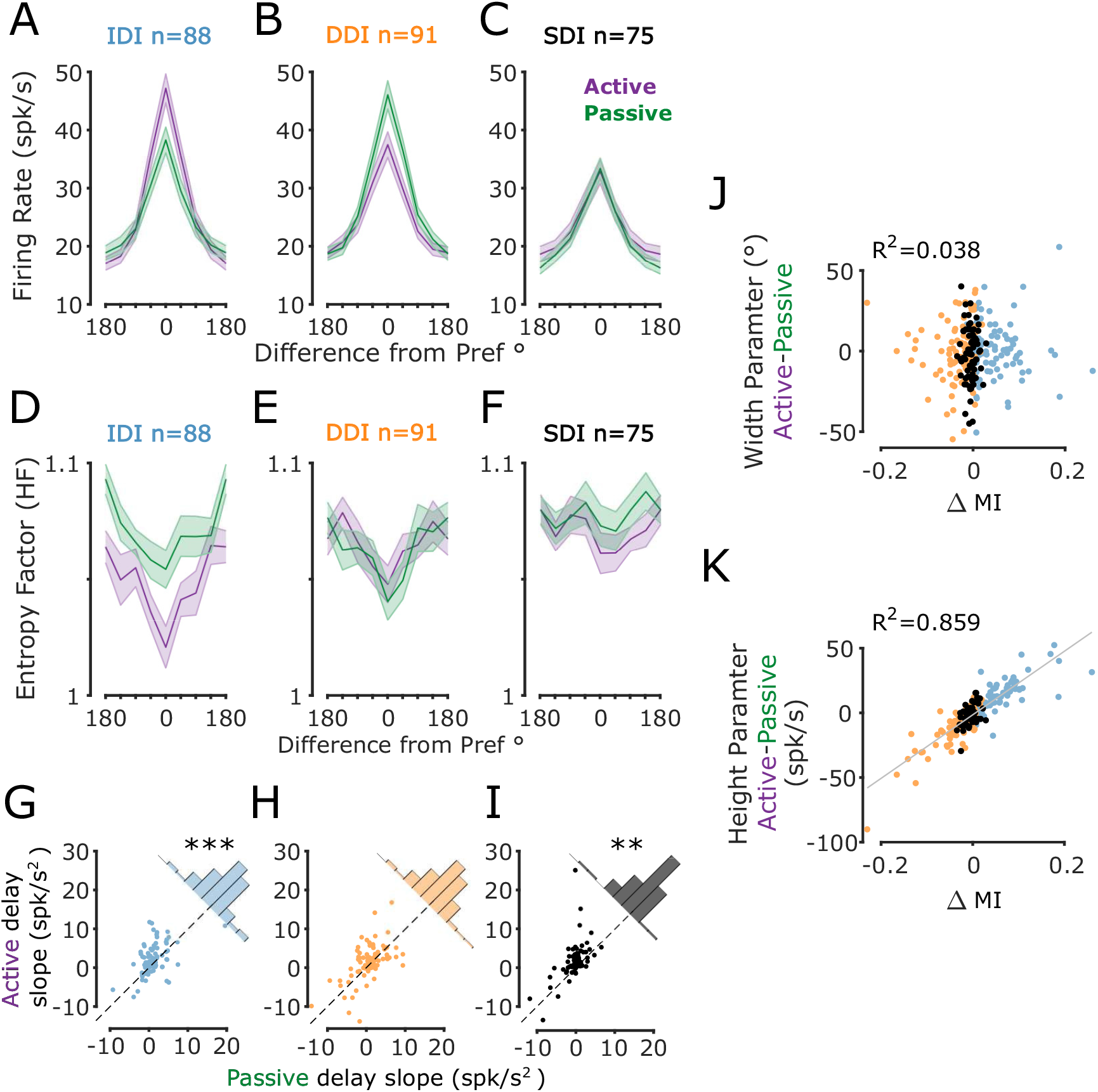
Effects of task demands differed depending on how informative a neuron was about task-relevant information. A.) Average raw tuning curves for IDI neurons during the active (purple) and passive (green) tasks. Both curves contain the same neurons. Error bars are ±*SEM* across neurons. B-C.) Same as A for DDI and SDI groups respectively. D.) Average trial-to-trial variability during S1 quantified with entropy factor (HF) (See Methods). E-F.) Same as panel D for DDI neurons and SDI groups respectively. G.) Slope of a linear fit to IDI neurons’ firing rates vs time during a 1s interval before S2 onset in the Active vs Passive tasks. There was a significantly higher ramping during the active task (***: *p* = 5.66 · 10^*-*5^, t-test). H-I.) Same as panel G for DDI (ns) and SDI (**: *p* = 0.0098) groups respectively. J.) Task effect of the width parameter of gaussian fits vs Δ*MI* during S1. For each cross validation fold, we fit a gaussian to the 50% of trials not used for the calculation of MI (i.e. the X and Y axes use different trials). We then averaged across folds, and subtracted across task to get a task effect of tuning width. We found no significant relationship between tuning width and Δ*MI* (*R*^2^ = 0.038,*p* = 0.552). K.) Same as J but for the height parameter of the gaussian fits (*R*^2^ = 0.859,*p* = 3.505 · 10^*-*75^).

Next, we sought to quantify how responses changed with task condition in terms of their direction tuning curves. Direction tuning curves can be conceived as bell-shaped functions with four parameters: amplitude, width, shift (preferred direction) and offset (baseline firing rate). To test whether the changes in tuning that we observed could be attributed to changes in one or more of these parameters, we fit Gaussian functions to the responses of each neuron as a function of stimulus direction, and then tested for an effect of task (‘active’ vs. ‘passive’) for each parameter with paired t-tests for each group of neurons (IDI, DDI, and SDI). For the IDI neurons, we found the amplitude parameter was significantly greater in the active task than in the passive task (change in amplitude for S1: −11.66 ± 1.34 sp/s [mean ± SEM], 86.09 ± 13.54%, *t*_87_ = −8.73, *p* = 2 · 10^−13^. For S2: 11.01 ± 1.55 sp/s, 98.11 ± 17.03%, *t*_86_ = −7.19, *p* = 2 · 10^−10^). All other parameters were not significantly different (all p’s > 0.07). In contrast, the DDI neurons showed lesser tuning curve amplitudes in the active task than in the passive task (change in amplitude for S1: −11.52 ± 1.66 sp/s, 28.46 ± 4.05%, *t*_89_ = 6.99, *p* = 5 · 10^−10^. For S2: −9.95 ± 1.44 sp/s, 28.58 ± 4.01%, *t*_89_ = 6.90, *p* = 7 · 10^−10^). All other parameters, including tuning width, were not significantly different (all p’s > 0.08). The SDI group showed slightly greater offset (baseline) values in the active task than in the passive task during S2 (*p* = 0.03), consistent with the main effect of ‘task’ for the ANOVA results reported above, but this was not robust to correction for multiple comparisons for the three groups of neurons tested. We further tested the relationship between tuning and Δ*MI* by considering the full distribution, void of any assumptions about clusters. We again calculated tuning curves for each neuron, using only the 50% of trials not used for the clustering analysis. This resulted in a distribution of parameters for each neuron across cross-validation folds, which we then averaged and plotted against Δ*MI* (Figure 4J-K). Again, only the amplitude (height) parameter could explain the differences in direction information (*R*^2^ = 0.859,*p* = 3.505·10^−75^). Taken together these results show that during the active task, neurons displayed a continuous, approximately multiplicative, modulation of the amplitude of their tuning functions. These task-based modulations likely contribute to the patterns of IDI, DDI, and SDI neurons observed previously.

### Trial-to-Trial Response Variability is Modulated by Task Demands

In addition to tuning, MI also takes into consideration trial-to-trial response variability. We reasoned that the task effects on MI may be partially due to altered variability of neural responses. To test this, we calculated the entropy of neural responses and normalized measurements relative to those of a simulated Poisson process matched for intensity (entropy factor (HF)).

We found that during the active task, IDI and SDI neurons showed an overall reduction in HF (3-way ANOVA: IDI: *F* = 60.24,*p* = 1.18 · 10^−14^; DDI: *F* = 0.02,*p* = 0.879; SDI: *F* = 10.67,*p* = 0.001 Figure 4D-F) compared to the activity recorded during the passive task. Additionally, all three classes of neurons had changes in trial-to-trial variability that was a function of motion direction (3-way ANOVA: IDI: *F* = 6.64, *p* = 8.14 · 10^−8^; DDI: *F* = 6.97, *p* = 2.98 · 10^−8^; SDI: *F* = 2.07, *p* = 0.044; Figure 4D-F). All three groups of neurons showed the lowest HF for their preferred direction and the variability increased progressively for less preferred directions. The reduction in variability and its dependence on motion direction was strongest in the IDI group, consistent with the improvement to both tuning and MI during the active task. While the DDI group also displayed an effect of motion direction on HF, unlike the IDI group, it did not show dependence on task demands. SDI neurons showed only a weak reduction in HF during the active task and a very modest dependence on motion direction.

### Task Demands Modulate Activity in MT During Non-stimulus Periods

We next turned our attention to the effects of varying task demands on MT activity during periods when motion stimuli were absent (pre-S1, during the delay between S1 and S2, and post-S2). Because no stimuli were present, effects during these intervals would be a likely reflection of cognitive processes (e.g., anticipation of S1 or maintenance of memoranda during the delay). Since the duration of the pre-S1 period was highly predictable (1000ms), a gradual increase (“ramping”) of firing rate during this period, is likely to reflect anticipation (see example neuron in Figure 1B). We quantified pre-S1 ramping as the slope of a linear fit to FR vs. time during a one second window before S1 onset. We found significant pre-S1 ramping only during the active task and only for the IDI group (T-test Passive minus Active slopes: *p* = 1.02 · 10^−4^).

Next, we examined the pattern of activity during the delay separating S1 and S2. Because the duration of this delay was also highly predictable (always 1.5 s) animals could anticipate the appearance of S2. Similar to the earlier report (Bisley et al. 2004), we found significant ramping in the delay activity during the active task and this ramping was most pronounced for the IDI neurons (Paired sample T-Test on delay slopes: *p* = 5.66 · 10^−5^) and SDI neurons (*p* = 0.0098; Figure 4G-I). This increased ramping in the active task may reflect additional task demands related to the preparation for S2, the comparison stage of the task.

### Effects of Task Demands on Comparison Signals

After S2, our active task required subjects to make a saccade to the right if the direction of the preceding S1 was the same (“same trial”), and a saccade to the left if the direction of the preceding S1 was different (“different trial”). It has previously been observed that MT neurons’ responses to physically identical S2 stimuli differed whether they came from “same” or “different” trials, which we termed a “comparison effect” (CE) (Lui and Pasternak 2011; Zaksas and Pasternak 2006). Here, we examined whether the activity during S2 showed CEs and asked: (1) are CEs different in the active versus the passive task (when no comparison is needed); and (2) if these effects differ across the three neuron groups (IDI, DDI, SDI)?

We calculated CE as the area under the receiver operating characteristics curve (AUC) within direction condition during S2 and the post-S2 interval, between “same” and “different” trials (Lui and Pasternak 2011). This metric quantifies the separation of the firing rate distributions for “same” and “different” trials, where a value of *AUC* = 0.5 means the two distributions are indistinguishable, values 0 ≤ *AUC <* 0.5 means the neuron fires more strongly to identical stimuli on “different” trials compared to “same” trials, and values 0.5 *< AUC* ≤ 1 means the neuron fires more strongly on “same” trials compared to “different” trials. On average across the population, we did not find significant CEs during S2 (Figure 5, Figure 6A,C), but we did find CEs during the post-S2 interval for some groups of neurons (Figure 5, Figure 6B,D). We next asked how these CEs were affected by changing task demands. We expected that task effects on cognitive signals may be independent from task effects on sensory processing, a result that would indicate a division of labor across MT neurons for sensory and cognitive processes: the effect of increasing task demands for a given MT neuron would be to either increase direction signaling, or increase comparison effects, but not both. Thus, neurons with stronger motion direction signals in the active task would not also increase comparison effects in the active task; whereas neurons that did not increase signaling of motion direction in the active task would be more likely to show increased comparison effects.

In line with this prediction, we found that DDI and SDI groups showed significantly stronger *S* > *D* CEs during the post-S2 period in the active task, the period immediately preceding the monkeys’ report (Figure 5C,E; DDI: Kolmogorov-Smirnov test: *p* = 4.46·10^−4^, SDI: Kolmogorov-Smirnov test: *p* = 0.0019 on average CE in window) and a higher proportion of neurons with significant CEs (Figure 6B; DDI: *χ*^2^ = 11.14, *p* = 8.43 · 10^−4^, SDI: *χ*^2^ = 14.23,*p* = 1.61 · 10^−4^). Interestingly, the DDI group showed weaker *D* > *S* signals in the active task compared to the passive task (Figure 5D; Kolmogorov-Smirnov test: *p* = 0.0011 on average CE in window). In contrast to the clear task effects on CEs for the DDI and SDI neurons, the IDI group of neurons showed no significant difference in CE magnitudes between the two tasks (Figure 5A-B). IDI neurons did show a trend for more prevalent *D* > *S* neurons, but this effect did not stand up to correction for multiple comparisons (Figure 6D, *χ*^2^ = 6.21,*p* = 0.013). When viewing the whole distribution of comparison effects vs direction information, it becomes increasingly clear that neurons with strong comparison effects convey less direction information (Figure 6E), despite being tuned for direction (Figure 4B-C). The higher proportion of *S* > *D* neurons in the DDI and SDI groups, along with their increased CE signaling, indicates that while the heightened task demands did not improve their stimulus encoding, these neurons may have been recruited to aid the comparison process.

**Figure 5.**
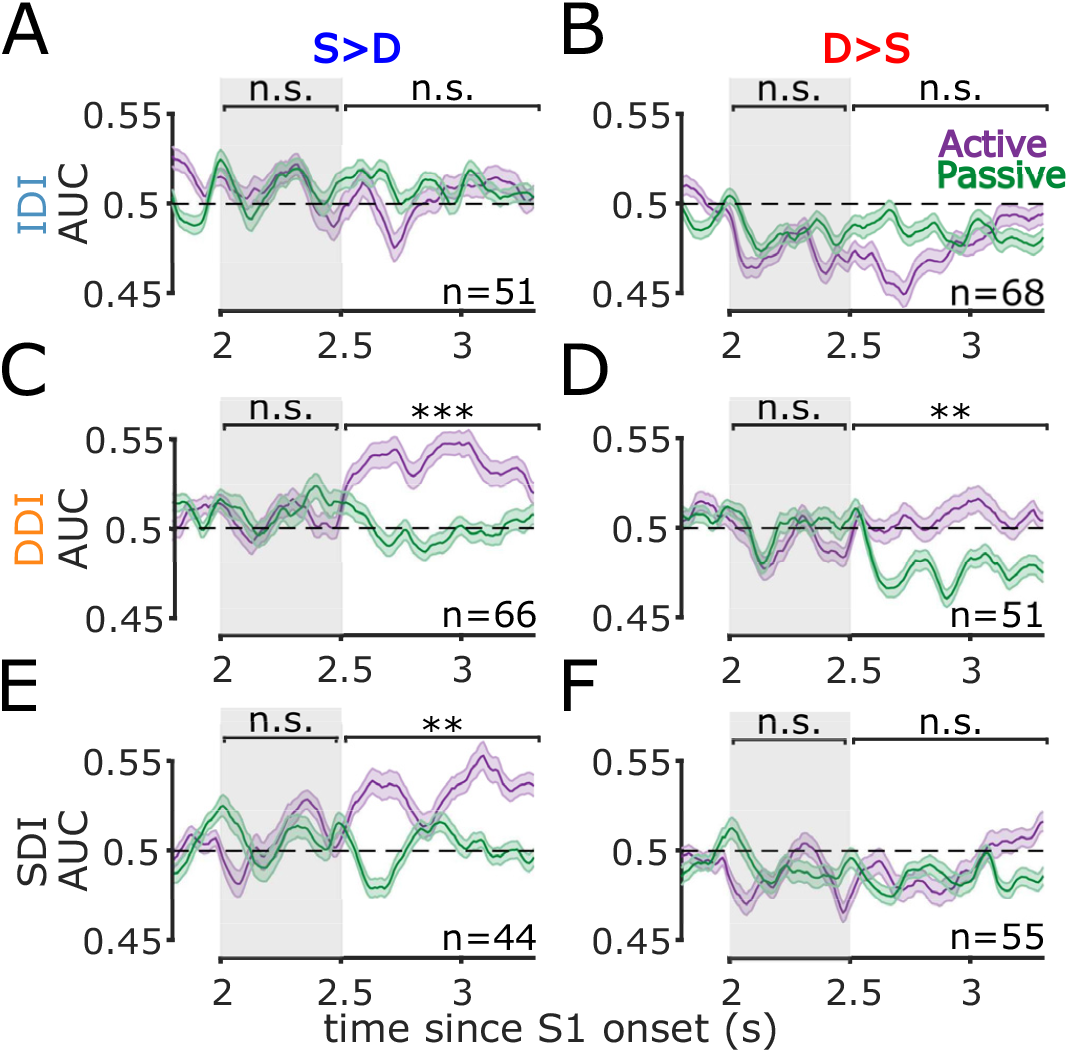
DDI and SDI neurons have different comparison effects across the active and passive tasks. A.) Time course of comparison effects (AUC between same and different trials within stimulus type) for the IDI neurons with significant *S* > *D* in either task. B.) Same as panel A but for IDI neurons with *D* > *S* signal. C-D.) Same as A-B but for DDI neurons. E-F.) Same as A-B but for SDI neurons. **: *p <* 0.01, ***: *p<* 0.001, n.s.: not significant t-test for the difference between tasks over the indicated interval. We did not find significant CEs or task effects during the S2 (gray shading) for any group of neurons, but we did see effects during the post-S2 interval for DDI neurons (C-D), and for SDI neurons with *S* > *D* CEs (E).

**Figure 6:**
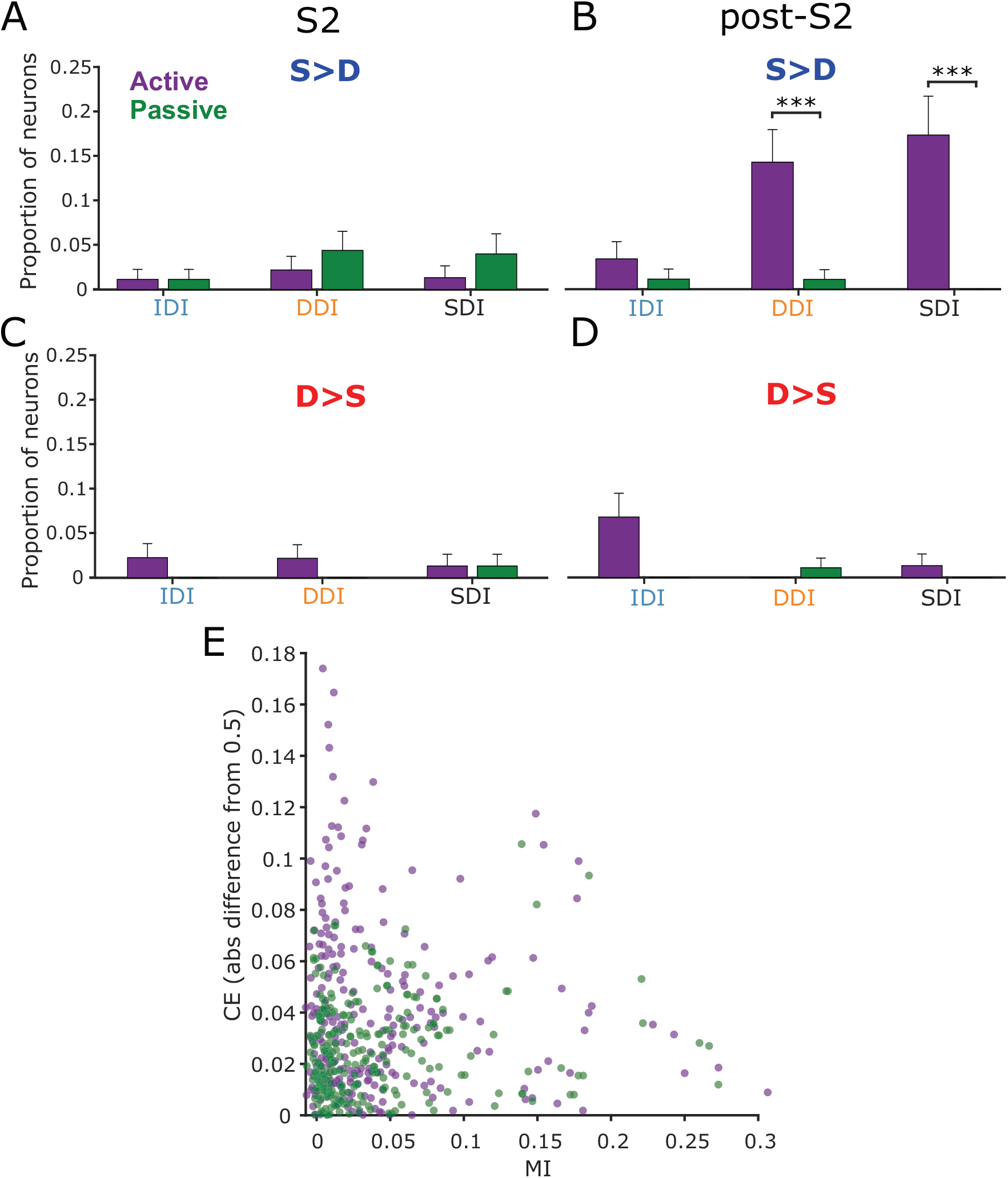
Proportion of neurons with significant comparison signals during active and passive tasks. A.) Proportion of neurons within each information group with significant *S* > *D* signal during the S2. B.) Same as panel A for the post-S2 interval. C) Same as in panel A for the *D* > *S* neurons. D) Same as in panel B for the *D* > *S* neurons. (***: *p <* 0.001, ^2^ test). E.) Comparison Effects (absolute difference from 0.5 AUC) vs mutual information for each task. Neurons with the largest CE in the active task also conveyed the least direction information

## Discussion

### Task Demands and Sensory Processing

We hypothesized that neurons in the primate Middle Temporal (MT) area behave differently when task demands increase in order to improve information processing for goal-oriented behavior. We tested this hypothesis by directly comparing responses to motion during two identically structured tasks: the active task that required monkeys to correctly report a decision for reward, and the passive task in which monkeys were rewarded on each trial without making the report. As the neurons were matched across tasks, we could directly compare how each neurons’ tuning and variability were impacted by task demands. Mutual information (MI) analysis used to quantify motion signals suggested heterogeneity of task effects distributed across roughly three groups of neurons: Increased Direction Information (IDI), Decreased Direction Information (DDI), and Similar Direction Information (SDI), each with different contributions to sensory and cognitive aspects of the motion comparison task. During the active direction comparison task, the IDI neurons showed a relative amplitude gain in direction tuning while the direction tuning of the DDI neurons became relatively weakened (Figure 4). This difference in tuning is likely to be one of the factors contributing to the change in the altered motion information during the two tasks. These effects on responses to motion are consistent with the classification of neurons based on changes to motion direction signaling in MI, but may not account for the full picture, as MI is also affected by trial-to-trial response variability.

The extent to which responses to motion and variability are each influenced by cognitive factors has been a subject of intense research (Renart and Machens 2014; Gilbert and Li 2013), and each has different implications for computation and neural mechanisms. We found a strong relationship between direction preference and trial-to-trial response variability, which did not depend on group (Figure 4D-F). Neurons had the largest reduction in trial-to-trial variability during presentations of their preferred stimulus, with weaker effects as direction diverged from preferred. This cannot be due to the increased firing rate for those directions, as entropy factor (HF) accounts for rate effects (see section Methods). Although the DDI neurons showed reduced tuning for direction they did not alter their trial-to-trial variability. Previous work has suggested that a substantial portion of trial-to-trial variability can be understood in terms of neurons’ susceptibility to changes in cognitive state (Ecker et al. 2016; Denfield et al. 2018); thus, one interpretation of this pattern of results is that these neurons decrease their contribution to sensory processing while maintaining their participation in more cognitive processes. The observed reduction in trial-to-trial stimulus response variability in IDI neurons with higher task demands is reminiscent of the finding of Mitchell, Sundberg, and Reynolds (2007), who found general reductions in Fano Factor in V4 responses when animals selectively attended to a stimulus compared to when that stimulus was not attended, as well as Hussar and Pasternak (2010) in the LPFC using a similar behavioral paradigm to the current experiment. The novel and more surprising result we found is that the strength of variability reduction depended on whether the stimulus was more or less preferred. The improved tuning for direction during the active task suggests that elevating task demands may increase reliability in the information transmission.

### Task Demands Affected Anticipatory Signals During the Delay

We found a significant difference in ramping of the delay activity recorded during the active (relative to passive) task (Figure 4G-I). Ramping of neural firing rates has traditionally been explored in the context of motor preparation (Ding 2015; Narayanan 2016), and decision making (Shadlen and Newsome 2001), where it has been typically linked to anticipation. Finding anticipatory increases in firing rate for sensory neurons in MT is consistent with previous reports (Bisley et al. 2004; Zaksas and Pasternak 2006) and is not likely explained during the pre-S2 interval of our tasks through either decision making or motor planning, as neither a correct decision nor appropriate motor plan can be made before first seeing S2. More likely, the ramping we observed is related to preparing cognitive and attentional resources for the perceptual phases of the task, which had highly predictable timing, as has also been found for LPFC neurons (Hussar and Pasternak 2010; Hussar and Pasternak 2013). This interpretation of ramping activity as preparing perceptual resources also aligns with our finding that pre-S1 ramping was restricted to IDI neurons, which enhance perceptual signaling in the active task. This line of thought has also been explored in the context of another nearby visual region, V4 (Snyder, Yu, and Smith 2018; Luck et al. 1997), where anticipatory ramping has been linked to interactions between V4 and prefrontal cortex (Snyder, Yu, and Smith 2021). Taken together, these results suggest anticipatory signals may originate in prefrontal cortex and influence visual cortex via top-down feedback.

### Task Demands and Comparison Effects

In addition to the more strictly sensory aspects of the task, the active task contains a comparison component which requires the subject not only to process the current stimulus direction (S2) but also retrieve the direction of the previously seen S1, perform the comparison and report it 1 second later. During the passive task, all the components of the task are present, but the animal is rewarded without having to report its decision. Previous work revealed that during the direction comparison task, similar to that used in the present study, responses of many MT neurons during S2 reflected the direction of the preceding S1, with some neurons showing stronger responses to S2 when its direction matched S1 (*S* > *D*) and other neurons showing stronger responses to S2 when its direction was different from S1 (*D* > *S*) (Lui and Pasternak 2011). Here, we examined if such signals, termed comparison effects, are also present when the task does not require the animals to make direction comparisons. We would expect that if CEs in MT are explained primarily by “passive” sensory effects such as adaptation/facilitation (Kohn and Movshon 2003), we should find similar effects in the active and passive tasks. In contrast, a difference in the magnitude or incidence of CEs between the active and passive tasks would indicate a greater role for memory-guided comparison processes underlying CEs. We found no difference in magnitude of comparison effects for IDI neurons at a population level, but a slight increase in incidence for significant *D* > *S* signals in this group. Taken together with the gain on tuning, it may be the case that this result reflects heightened adaptation in the active task, which would be consistent with a primarily sensory explanation of CEs for these neurons. However, we found significantly stronger (Figure 5) and more prevalent (Figure 6) CEs for the DDI and SDI groups, suggesting a greater cognitive role for CEs for these cells. These cognitive effects being isolated to neurons that became worse at signaling motion direction when that information became task-relevant further supports a division of labor in area MT for completing the task. That is, these neurons shed direction signaling in favor of increased CEs when active comparison was required for the task.

We found that comparison effects emerged most clearly during the period after the offset of S2 and before the execution of the response saccade (the ‘post-S2’ interval; Figure 5). This underscores the non-sensory, cognitive nature of these effects (since the sensory stimulus was absent when effects were strongest) and reduces the potential that response adaptation or repetition suppression could explain D>S comparison effects (Lui and Pasternak 2011; Kohn and Movshon 2003). This relatively late time-course of comparison effects in MT differs from earlier reports, which found comparison effects emerging shortly after S2-onset (Lui and Pasternak 2011; Zaksas and Pasternak 2006). One reasonable explanation for this difference in the timing of comparison effects across studies is the difference in timing between the onset of S2 and the animals’ reports. Earlier studies required the animals to respond immediately following the offset of the S2, whereas in the current study animals must withhold responding until after the post-S2 delay. Thus, if the timing of comparison effects was more tightly related to the animals’ reports than to the time of S2 onset, then that would be consistent with the results across all studies of comparison effects in MT and further underscores the cognitive role of comparison signals. Finally, in earlier studies decisions were reported with a manual button press (Lui and Pasternak 2011; Zaksas and Pasternak 2006), whereas the current study required animals to report with a saccade, therefore CEs are independent of the mode of decision reporting, reinforcing the notion that comparison effects are related to cognitive decisions rather than the formation of particular motor plans.

## Conclusion

We investigated the effects of task demands on information processing in area MT of the macaque. We found a diversity of neurons that were affected in different ways in the active compared to the passive task. Neurons that increased their signaling of task-relevant sensory information (IDI neurons) did not show changes in comparison effects, which reflect more cognitive memory and comparison processes, whereas neurons that either decreased (DDI) or did not change (SDI) direction information in the active task showed changes in their comparison signals. These results suggest an distributed allocation of labor of cognitive and perceptual processes across a population of neurons in MT and highlight the role of MT populations in all phases of memory-guided comparisons of visual motion direction.

## Acknowledgements

This work was supported by NIH grant R01EY011749 (T.P. and A.C.S.); NIH grant R00EY025768 (A.C.S.); a Sloan Research Fellowship (A.C.S.), NARSAD Young Investigator Award (A.C.S.), as well as the Spanish State Research Agency together with the European Regional Development Fund (grants RYC-2015-17236 (K.W.), BFU2017-86026-R (K.W.), PID2020-112838RB-I00 (K.W), and, through the Severo Ochoa and María de Maeztu Program for Centers and Units of Excellence in R&D, CEX2020-001084-M). KW thanks CERCA Programme/Generalitat de Catalunya for institutional support.

## Author Contributions

This project was conceptualized by T.P.. Data acquisition was done by T.P.. Data analysis was done by H.S., with guidance from A.S. and K.W. Original draft of manuscript was written by H.S. with A.S., K.W., and T.P. providing editorial and revision feedback.

## List of Abbreviations

DVA: degrees of visual angle
FR: firing rate
HF: entropy factor
LPFC: lateral prefrontal cortex
MI: mutual information
MT: Middle Temporal
PCA: principal component analysis
RF: receptive field

---

**Table 1:** Task effects summary. Significance of task effects (p value) separated by information clusters. Statistics for Tuning and entropy (HF) were a three-way anova between task, motion direction, and stimulus (S1 vs. S2). Delay ramping was compared using t-tests on the slopes (Passive minus active: (4 G-I).

| Class | Tuning |  |  | HF |  |  | Delay<br>Ramping |
| --- | --- | --- | --- | --- | --- | --- | --- |
|  | Task | Dir. | Stim. | Task | Dir. | Stim. |  |
| **IDI** | $9.54 \cdot 10^{-5}$ | $5.47 \cdot 10^{-126}$ | 0.238 | $1.18 \cdot 10^{-14}$ | $8.14 \cdot 10^{-8}$ | 0.018 | $5.66 \cdot 10^{-5}$ |
| **DDI** | $1.97 \cdot 10^{-5}$ | $1.49 \cdot 10^{-140}$ | 0.187 | 0.879 | $2.98 \cdot 10^{-8}$ | 0.011 | 0.557 |
| **SDI** | 0.005 | $2.70 \cdot 10^{-66}$ | 0.531 | 0.001 | 0.044 | 0.021 | 0.010 |

