## Supplemental Figures for "Altered Task Demands Lead to a Division of Labor for Sensory and Cognitive Processing in the Middle Temporal Area"

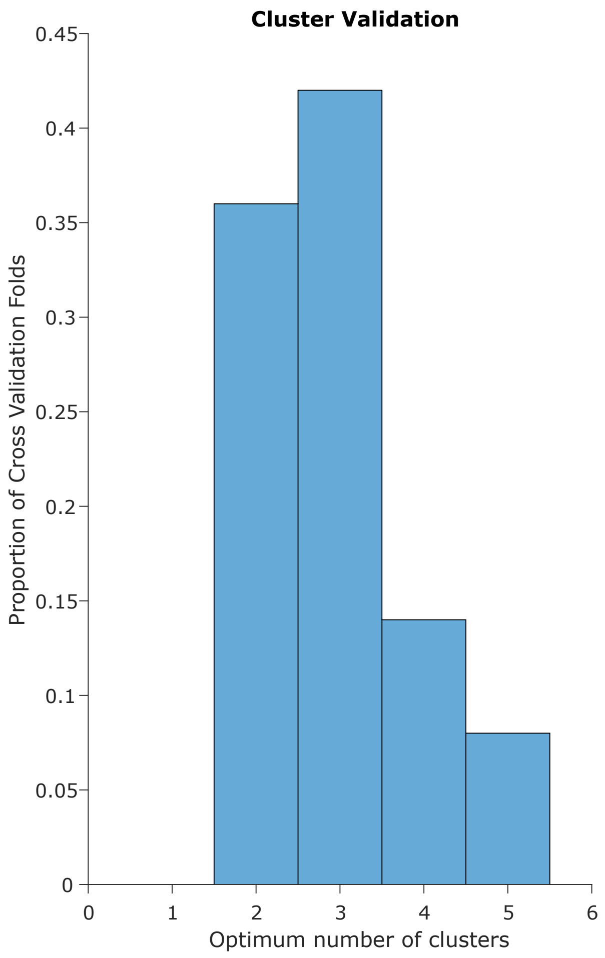


Supplemental Figure 1: Cross validation of clustering algorithm. We took 50% of trials for each neuron, calculated MI, ran the clustering algorithm, and minimized AIC to pick the optimum number of clusters. After repeating this process 100 times, we got a distribution of models that supports 3 clusters as optimal for our data.

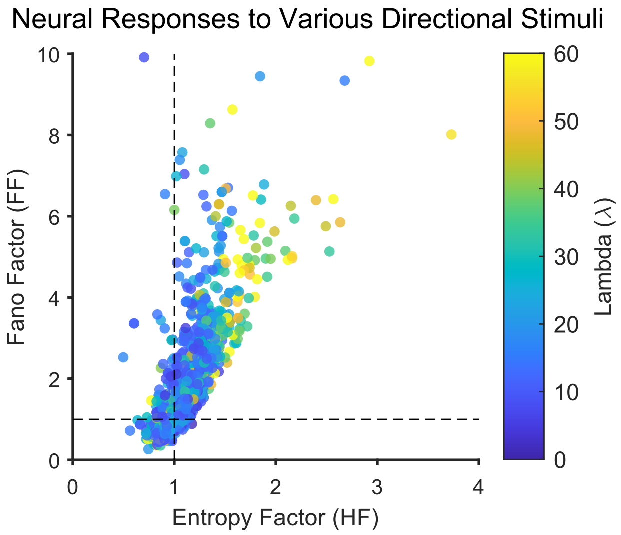


Supplemental Figure 2: Relationship between Entropy Factor (HF) and Fano Factor (FF). We took the neural responses for our 8 directions of motion (S1), for each neuron in our sample, and plotted FF vs HF. Dotted lines are Poisson processes. Neural responses with a larger lambda parameter (average firing rate) result in a roughly linear relationship between the measures. When neural responses are low, FF overestimates the variability (darker blue dots are highly super-Poisson according to FF, but roughly Poisson according to HF)

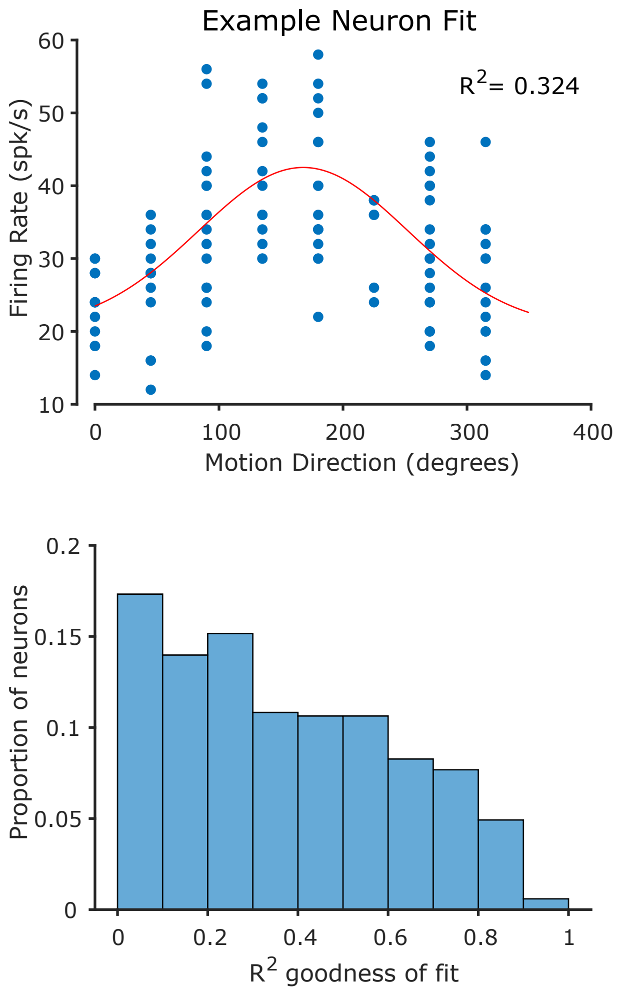


Supplemental Figure 3: Example and overview of tuning goodness of fit ($R^2$). A Gaussian fit to one example neuron during one bootstrap fold. The full distribution of $R^2$ for our sample of neurons.

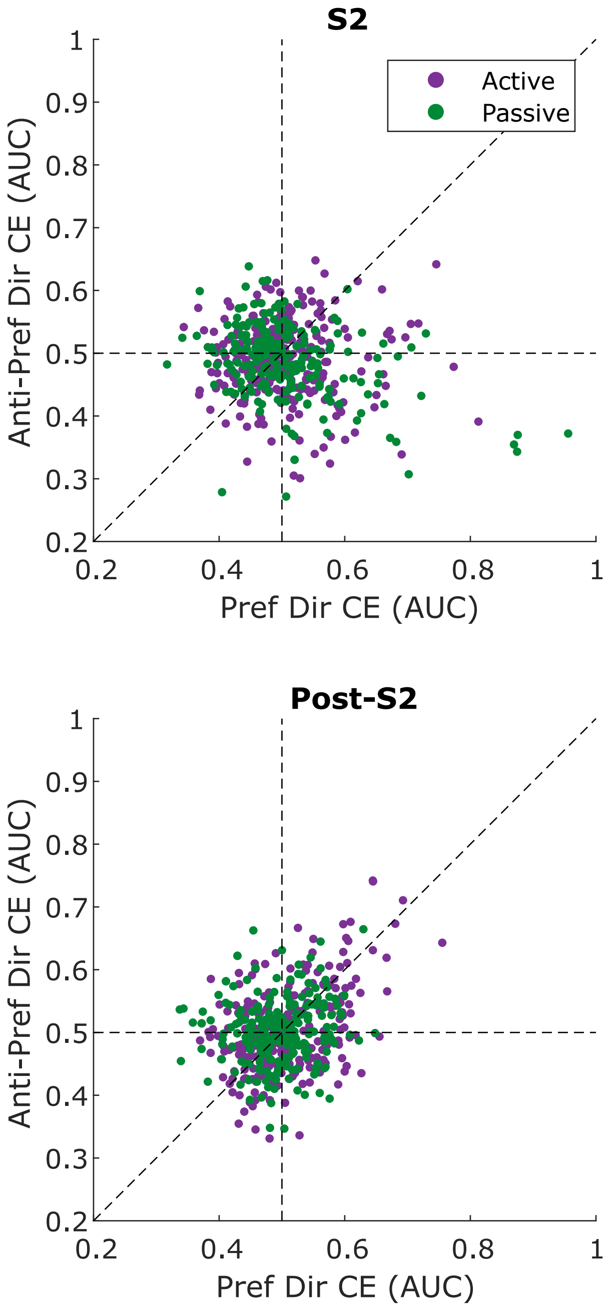


Supplemental Figure 4: Average Comparison Effect (CE) during the two windows of interest for neurons' preferred direction and 180° off preferred (anti-preferred).
